# First report of antiviral activity of carbon nanofibers: Enhancement of the viral inhibition capacity of calcium alginate films

**DOI:** 10.1101/2020.08.18.255646

**Authors:** Isaías Sanmartín-Santos, Sofía Gandía-Llop, Ángel Serrano-Aroca

## Abstract

The World Health Organization has called for new effective and affordable alternative antiviral materials for the prevention and treatment of viral infections. In this regard, calcium alginate has previously shown to possesses antiviral activity against the enveloped double-stranded DNA herpes simplex virus type 1. However, non-enveloped viruses are more resistant to inactivation than enveloped ones. Thus, the viral inhibition capacity of calcium alginate and the effect of adding a minuscule amount of carbon nanomaterials (0.1% *w/w*) have been explored here against a non-enveloped double-stranded DNA virus model for the first time. The results of this study showed that neat calcium alginate films are able to inactivate this type of non-enveloped virus and that including that extremely low percentage of carbon nanofibers significantly enhanced its viral inhibition from ~55.6% to 96.33%. This is the first published study to demonstrate CNFs’ antiviral activity. However, adding this small percentage of graphene oxide did not improve the antiviral activity of calcium alginate, although both composite biomaterials possess antiviral and other outstanding properties very promising for biomedical applications.

## 1. Introduction

Sodium alginate (SA) has been authorized by the US Food and Drug Administration for human biomedical applications due to its excellent properties such as biodegradability, renewability, cost-effectiveness, non-toxicity and biocompatibility [1]. This biopolymer can be cross-linked with Ca^2+^ cations to form hydrogels [2] with many physical properties that can be enhanced by adding very small amounts of carbon nanomaterials (CNMs) such as graphene oxide (GO) [3–5] or carbon nanofibers (CNFs) [6,7]. These nanocomposites have similar biological properties to neat calcium alginate in terms of cell adhesion [8] and non-cytotoxicity [9,10] and GO is known to be antibacterial [10]. We have recently report for the first time that pure CNFs or added to calcium alginate, have antibacterial properties that can be exploited to eradicate multidrug resistant pathogens [9]. In general, non-enveloped viruses are more resistant to inactivation than enveloped virus [11]. Calcium alginate-based materials have shown antiviral activity against the enveloped double-stranded DNA herpes simplex virus type 1 (HSV-1)[12]. However, calcium alginate’s antiviral activity and the effect of incorporating a minuscule amount (0.1% *w/w*) of carbon nanofibers or graphene oxide has never been studied before using a non-enveloped double-stranded DNA viral model [13]. Based on these previously published antiviral results [12], we hypothesized that calcium alginate could exhibit antiviral activity against this different viral model belonging to Group I of the Baltimore Classification [14] like HSV-1. Since graphene oxide has been shown to be antiviral against DNA and RNA viruses such as the pseudorabies virus and porcine epidemic diarrhea virus [15], we also considered that incorporating these two carbon nanomaterials would enhance their antiviral action. As far as we know, the antiviral activity of pure CNFs or added to calcium alginate has never been studied before.

## 2. Materials and Methods

### 2.1. Materials

Sodium alginate (Panreac AppliChem, Darmstadt, Germany), calcium chloride (anhydrous, granular, ≤7.0 mm, ≥93.0%, Sigma-Aldrich, Saint Louis, Missouri, USA), graphene oxide (Ref: 796034, powder, 15-20 sheets, 4-10% edge-oxidized, Sigma-Aldrich, Saint Louis, Missouri, USA) and carbon nanofibers (Ref: 13/0248, Graphenano, Yecla, Spain) were used as received.

### 2.2. Synthesis

Alginate nanocomposite films of approximately 0.25 g were prepared with a composition of 99.9% *w/w* of SA and 0.1% *w/w* of GO or CNFs following a recently reported new engineering route to produce more homogenous alginate-based composites with enhanced physical properties [4]. GO/SA or CNFs/SA was mixed in 22 ml of distilled water by magnetic stirring for 1 hour at room temperature (26±0.5°C), after which another aqueous solution containing 6% (with respect to the SA mass) of CaCl2 in 10 ml of distilled water was mixed with the GO/SA or CNFs/SA aqueous solution for 10 more minutes. Thin films were produced in Petri dishes after 24 h of drying at 37±0.5°C in an oven by solvent evaporation. The films were cross-linked by immersion in an aqueous calcium chloride solution (2% *w/v*) for 2 hours and after rinsing with distilled water were vacuum dried at 60°C±0.5°C. The calcium alginate films without CNMs were produced by the same chemical procedure. These films will be referred to hereinafter as *Alginate, GO0.1%* and *CNFs0.1% films*.

### 2.3. Characterization

#### 2.3.1. Biopolymer characterization

The sodium alginate used was characterized by nuclear magnetic resonance (NMR), high-performance anion-exchange chromatography with pulsed amperometric detection (HPAEC-PAD) and size exclusion chromatography with multi angle light scattering detection (SEC-MALS) by the NOBIPOL research group at the Norwegian University of Science and Technology (NTNU).

#### 2.3.2. Electron microscopy and Raman spectroscopy

The GO nanosheets and CNFs were examined by high-resolution transmission electron microscopy (HR-TEM) in a JEM 2100F (JEOL, Japan) 200 kV electron microscope with energy-disperse X-ray spectroscopy (EDS) at 20 kV. The sample preparation consisted of dispersing a very small amount of CNMs in dichloromethane in an ultrasound bath for ten minutes and then drying at ambient temperature before HR-TEM observation. A JEM-1010 (JEOL, Japan) 100 kV transmission electron microscope (TEM) was utilized to observe the CNFs and GO nanomaterials incorporated into the calcium alginate films. Ultrathin samples with 60 nm sections were prepared on a Leica Ultracut UC6 ultramicrotome (Leica Mikrosysteme GmbH, Austria) and a Diatome diamond knife (Diatome Ltd., Switzerland). The specimens were placed on TEM grids (300 mesh) coated in carbon film.

Raman spectroscopy was performed from 1000 to 3000 cm−^1^ in a Renishaw inVia confocal micro-Raman apparatus at 600 L·mm−^1^ grating equipped with an argon ion laser at 633 nm edge (10% power), ×20 lens, and a Renishaw CCD camera detector. The pure CNMs and or added to the calcium alginate films were placed onto glass disks for direct Raman analysis.

#### 2.3.3. Antiviral activity tests

The *Escherichia coli* B, living, bacteriophage host (Reference: 12-4300) and the coliphage T4r (Reference: 12-4335) from Carolina (Burlington, North Carolina, USA) were used for the antiviral tests. The sample films of alginate, GO0.1% and CNFs0.1% were cut into 1 mm diameter discs and sterilized by washing the films for 30 minutes in a beaker with 50 µl of 70% ethanol under magnetic stirring at 450 r.p.m. with fresh ethanol every 10 minutes. The discs were then dried at room temperature and exposed to ultraviolet light for 1 hour per side. The films’ antiviral activity was analyzed by a “contact test” of the bacteriophages with the discs in 96-well plates for 0, 18 and 48 hours. Several 1/10 serial dilutions from the original concentrate phage stock of about 1×10^9^ plaque forming units (PFU) per ml were tested and the 10^−5^ dilution was chosen from this stock as optimal for our experiments. 50 µl of this dilution contained approximately 500 PFU. The dry film disc was placed in a precision balance and drops of phosphate buffer saline (PBS) solution were added with a volumetric micropipette until the film was saturated without excess liquid. The correct volume for the saturation point (44 µl of liquid per disc) was determined gravimetrically by the increase in weight. For the “contact test”, 44 µl of the 10^−5^ phage dilution was added to each disc and the 96-well plate was subsequently sealed with plastic film to prevent evaporation during the assay. After each test, the wells were filled with 500µl of PBS for washing. The washes were made by adding the buffer and performing 20 resuspensions across the surface of the disc with the help of the volumetric micropipette, after which the discs were extracted using a sterile forceps and stored with their corresponding liquid in 15 µl tubes and sonicated for 5 minutes at an amplitude of 10. The tubes were then vortexed for 1 minute and all the discs of the samples of contact at the different times were titrated. Six discs of each material were used to provide reproducible results at the three different contact times.

The infectious phage particles were determined by the “double layer” assay method. Briefly, the phage aliquots extracted from the disc were serial diluted (1/10 dilution factor) in PBS. An aliquot of 100 µl of each dilution was mixed with 150 µl of bacterial culture and 3 ml of the “top agar” (0.75% w/w agar in LB medium) preheated in a bath incubator to 45°C. The *E. coli* B culture was grown overnight until reaching an optical density of 1.0 at 600nm. The samples were gently shaken and poured over Petri dishes with the “bottom agar” (1.5% *w/w* agar in LB medium). The viable PFU viral particles were calculated from the plaques appearing on the soft agar surface stained with crystal violet.

#### 2.3.4. Statistical analysis

The statistical analyses were performed by ANOVA followed by Tukey’s posthoc test (***p > 0.001) on GraphPad Prism 6 software.

## 3. Results and Discussion

### 3.1. Characterization of the sodium alginate

The results of the sodium alginate characterization by NMR, SEC-MALS and HPAEC-PAD showed: a guluronic acid content of 43% and an alginate content consisting of guluronic acid in blocks of dimers and trimmers at 27 and 23% respectively, a weight-average (*Mw*) and a number-average (*Mn*) molecular weight of 379.5±9.5 and 170.7±3.1KDa, respectively.

### 3.2. Morphology, elementary composition and Raman analysis

The GO and CNFs in the form of powder were observed by high-resolution transmission electron microscopy (HR-TEM) and showed a morphology of irregular nanometric sheets and micrometer-length hollow fibers with a wide range of nanometric diameters, respectively (see Figure 1 (a&b)). The EDS results of the CNFs and GO showed C/O ratios of 31.3 and 15.4, respectively. The CNMs Raman spectra were also analyzed in that study and showed an ID/IG ratio of 0.92 for the hydrophilic 2D material (GO nanosheets) and 1.51 for the hydrophobic 1D filamentous CNFs, due to their higher degree of molecular disorder [16]. The TEM micrographs (Figure 1(c&b)) of the nanocomposites shows that the GO nanosheets or CNFs (dark phase) are embedded and randomly distributed in the alginate polymer matrix (clear phase).

**Figure 1.**
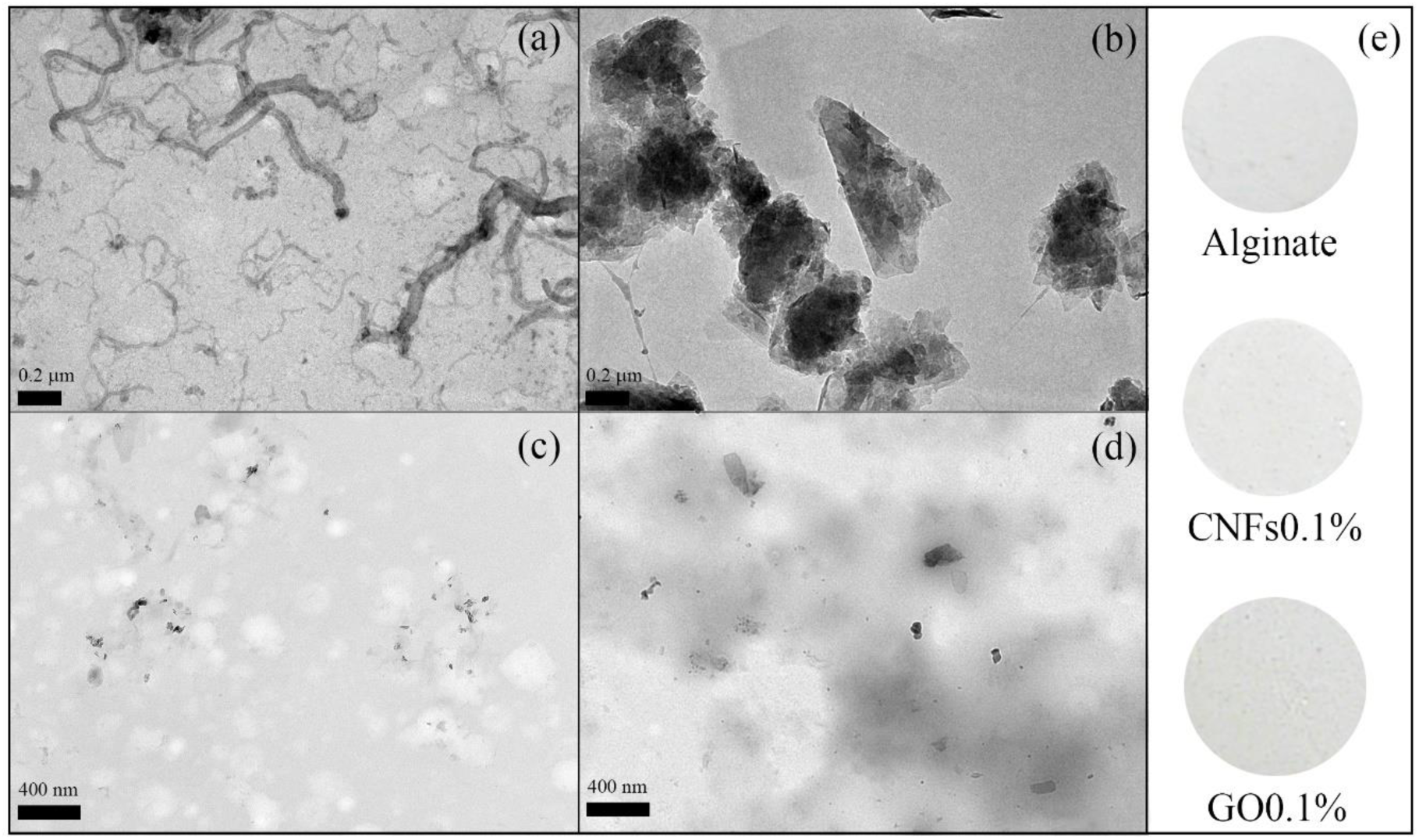
HR-TEM of carbon nanofibers (a), graphene oxide nanosheets (b), calcium alginate films with 0.1% *w/w* of carbon nanofibers (c), calcium alginate films with 0.1% *w/w* of graphene oxide (d) and digital photograph of the films (e).

Figure 1(e) shows that these antiviral nanocomposite films produced a negligible reduction of transparency, in good agreement with previous studies performed by our research group neither by adding carbon nanofibers [7] nor by adding graphene oxide nanosheets [10].

### 3.3. Antiviral properties

This was the first study carried out on the antiviral properties of calcium alginate with a small content of carbon nanofibers and graphene oxide. Figure 2 shows the plaque forming units measured in the antiviral tests after 0, 18 and 48 hours of contact with coliphage T4.

**Figure 2.**
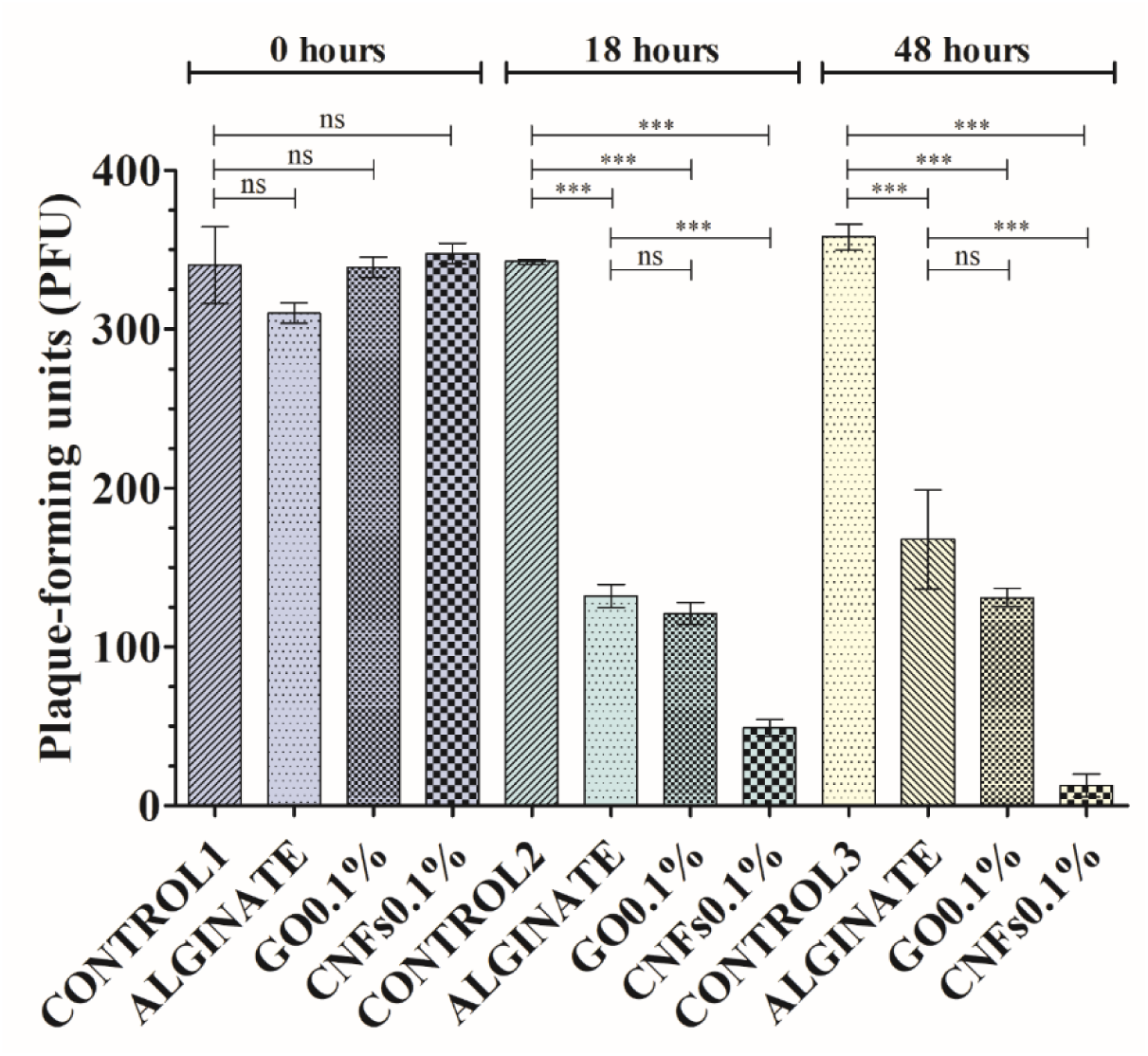
Antiviral activity results of control, alginate films, alginate/graphene oxide composite films (GO0.1%) and alginate/carbon nanofibers composite films (CNFs0.1%) after 0, 18 and 48 hours of contact with the coliphage T4 viral model.***p > 0.001; ns: not significant.

As expected, there was no statistically significant antiviral activity in the samples at 0 hours, indicating that the virus is not retained in the sample films after the sonication-vortex treatment, and that they needed some contact time to be inactivated. However, after 18 hours contact with coliphage T4 the viable phage counts were reduced by ~56-55.6% (see Figures 2 and 3).

**Figure 3.**
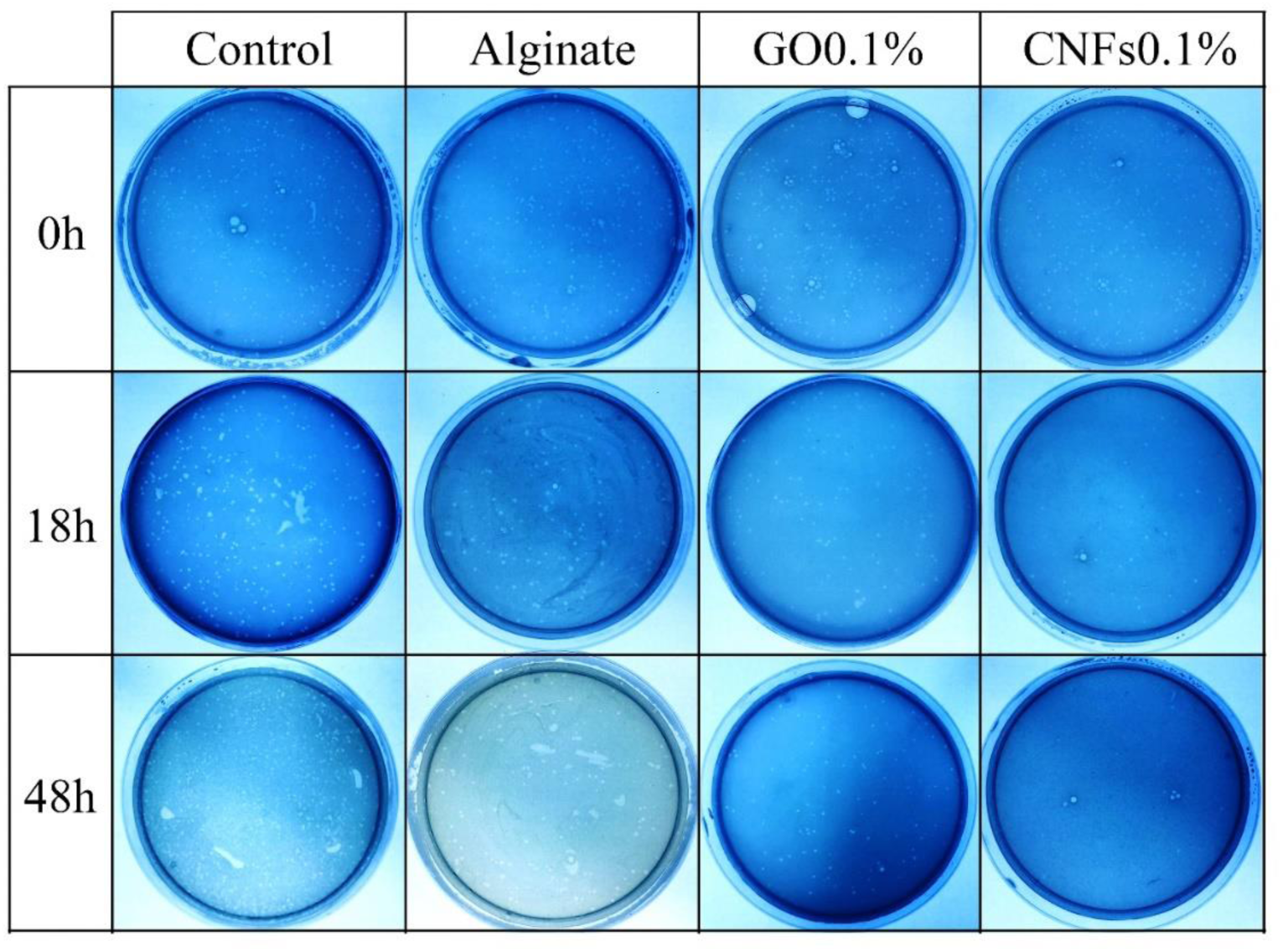
Representative titration plaques of coliphage T4: antiviral tests of alginate films, alginate/graphene oxide films (GO0.1%) and alginate/carbon nanofibers films (CNFs0.1%) at 0, 18 and 48 hours of contact with the virus.

Neither did calcium alginate’s antiviral action improve after a longer exposure period (48 hours), or with the incorporation of graphene oxide (see Figures 2 and 3). However, the calcium alginate films with a 0.1% carbon nanofiber content achieved viral inactivation of ~85.51 after 18 hours and up to 96.33% after 48 hours, showing that CNFs are strong antiviral agents. Even though viruses and bacteria are completely different, these results are in good agreement with previous antibacterial tests on calcium alginate films with the same GO[10] or CNFs[9] contents (0.1% *w/w*), which also found that the antimicrobial activity only improved with the incorporation of CNFs. Furthermore, these composite antiviral materials slightly reduce their transparency [7,10] and thus have an added advantage for use in transparent biomedical applications such as ophthalmology and odontology.

## 4. Conclusions

In this study we showed that, unlike graphene oxide, a minuscule amount of carbon nanofibers (0.1% *w/w*) can significantly enhance the antiviral activity of calcium alginate and also, for the first time in the literature, that CNFs have antiviral properties. Both alginate-based nanocomposites can cause viral inhibition, besides having enhanced physical and biological properties which render them potentially effective for use in antimicrobial biomedical applications.

## Acknowledgments

The authors are grateful to the NOBIPOL Group at the Norwegian University of Science and Technology (NTNU) for the characterization of the sodium alginate used in this study. They also wish to acknowledge the financial support through Grants 2019-231-001UCV and 2020-231-001UCV (awarded to Á.S-A) from the *Fundación Universidad Católica de Valencia San Vicente Mártir*.

## Data availability

The raw/processed data required to reproduce these findings cannot be shared at this time as the data also forms part of an ongoing study.

